# Conversion of graded presynaptic climbing fiber activity into graded postsynaptic Ca^2+^ signals by Purkinje cell dendrites

**DOI:** 10.1101/423491

**Authors:** Michael A. Gaffield, Jason M. Christie

## Abstract

The brain must make sense of external stimuli to generate relevant behavior. We used a combination of *in vivo* approaches to investigate how the cerebellum processes sensory-related information. We found that the inferior olive encodes contexts of sensory-associated external cues in a graded manner, apparent in the presynaptic activity of their axonal projections in the cerebellar cortex. Further, individual climbing fibers were broadly responsive to different sensory modalities but relayed sensory-related information to the cortex in a lobule-dependent manner. Purkinje cell dendrites faithfully transformed this climbing fiber activity into dendrite-wide Ca^2+^ signals without a direct contribution from the mossy fiber pathway. These results demonstrate that the size of climbing fiber-evoked Ca^2+^ signals in Purkinje cell dendrites is largely determined by the firing level of climbing fibers. This coding scheme emphasizes the overwhelming role of the inferior olive in generating salient signals useful for instructing plasticity and learning.

## Introduction

The cerebellum is thought to use sensory prediction errors to guide learning by plastic alteration of its circuitry. Climbing fibers (CFs) that arise from the inferior olive are well-suited to transmit sensory prediction errors to Purkinje cells (PCs), the sole output neurons of the cerebellar cortex. This is because each CF stimulation reliably initiates both a PC somatic complex spike and a dendrite-wide burst of Ca^2+^ potentials (Llinas and Sugimori, 1980). PCs encode descriptive information regarding sensory prediction errors by interpreting the population dynamics of these CF inputs (Brown and Raman, 2018; Herzfeld et al., 2018), as well as in the variance of evoked somatic and dendritic responses that distinguish behaviorally relevant CF events from spontaneous activity (Kitamura and Hausser, 2011; Najafi et al., 2014a, b; Yang and Lisberger, 2014). Such variability in discrete CF-evoked Ca^2+^ events may promote efficient coding of Ca^2+^-induced plasticity and learning in PC dendrites (Najafi and Medina, 2013; Rowan et al., 2018). However, the neural circuit mechanisms that support graded coding of CF-mediated signals are not fully resolved.

Olivary projection neurons fire bursts of spikes whose duration is influenced by the intrinsic dynamics of subthreshold oscillatory activity (Crill, 1970; De Gruijl et al., 2012; Leznik and Llinas, 2005). Therefore, the inferior olive is poised to directly convey processed information pertaining to external cues to the cerebellar cortex through variable-duration CF burst firing, particularly if their activity is subject to sensory amplification (Maruta et al., 2007). Yet, mossy fibers also relay sensorimotor information to PCs by way of granule cells. Through direct excitation and feedforward inhibition, granule cell activity can contribute to and/or modulate the CF-mediated dendritic Ca^2+^ response (Callaway et al., 1995; Wang et al., 2000).

In this study, we examined how PC dendrites process sensory-related information in the lateral cerebellum using a combination of *in vivo* Ca^2+^ imaging and optogenetics. We report that individual PC dendrites are reliably responsive to different modalities of sensory stimuli and remarkably sensitive to the presynaptic activity level of CFs. The absence of any additional, residual Ca^2+^ response from non-olivary inputs supports the conclusion that PCs faithfully convert sensory-enhanced CF excitation into graded, dendrite-wide Ca^2+^ signals. Thus, CFs provide near-exclusive sources of the instructive signals thought to be necessary for plasticity and learning in the cerebellum.

## Results

### Sensory stimuli trigger enhanced Ca^2+^ signals in PC dendrites

We used two-photon laser scanning microscopy (2pLSM) to measure Ca^2+^ signals in the dendrites of GCaMP6f-expressing PCs located in the lateral posterior cerebellum of awake, head-fixed mice (***Figure 1A***). Dendritic spikes, attributable to the activity of presynaptic CFs, were resolved as rapidly rising Ca^2+^ events (Ozden et al., 2008; Schultz et al., 2009); their continuous occurrence reflecting the regular spontaneous output of the inferior olive (1.01 ± 0.05 Hz). Next, we used a variety of sensory stimuli to activate the cerebellum and examine for evoked responses in PCs (***Figure 1B***). For this, we used somatosensory (airpuff to the ipsilateral whiskers [18 psi; 150 ms]), auditory (pure 12 kHz tone played at ~85 dB for 150 ms), and visual (light flash directed to the ipsilateral eye [λ = 473 nm; 25 μW; 150 ms]) stimuli. We presented each sensory modality in a block of 15 closely spaced trials (0.5 Hz), randomly switching between stimulus types from one block to the next to prevent habituation.

**Figure 1.**
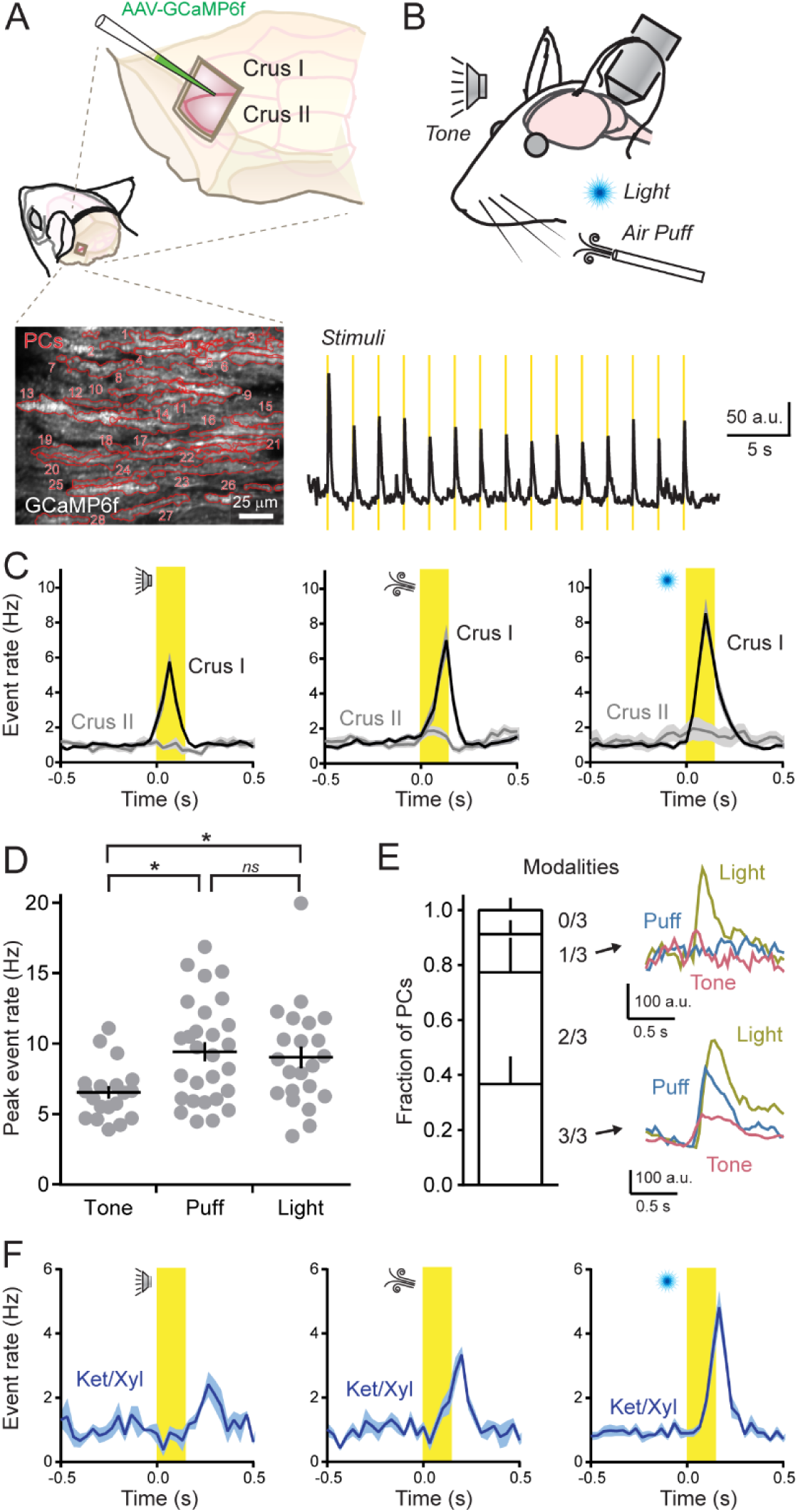
PCs in Crus I but not Crus II are broadly responsive to different sensory modalities. (**A**) Cranial windows were positioned over the left lateral cerebellum where PCs expressed GCaMP6f as shown in the fluorescence image. Red lines demarcate the dendritic contours of individual cells (n = 28). (**B**) Top: Sensory stimuli were presented to mice while measuring neural activity measurements. Bottom: Average ensemble Ca^2+^ response across PC dendrites (n = 20 trial blocks). Stimulus timing is shown in yellow. (**C**) Trail-averaged dendritic Ca^2+^ event rates for each sensory modality (i.e., tone, puff, light) across all PCs, compiled from measurements obtained from two different cerebellar lobules (n = 5-28 trial blocks from 6–10 mice per condition). (**D**) The peak rate of evoked Ca^2+^ events in Crus I PCs for each sensory modality across awake mice (each dot represents the average of a block of stimulus trials; 6-10 mice for each condition). Data are mean ± SEM with p = 0.01 for tone-puff comparison, p = 0.04 for tone-light comparison, and p = 0.90 for puff-light comparison; ANOVA with Tukey’s multiple comparison test. (**E**) Left: Fraction of identified PCs in Crus I, categorized based on their responsiveness to presentation of different sensory modalities (n = 141 cells from 3 mice). Right: Average dendritic Ca^2+^ transients from two different PCs that were either selectively (top) or broadly (bottom) responsive to modality types. (**F**) Dendritic Ca^2+^ event rates, measured in Crus I PCs, in response to sensory stimuli while the animal was under anesthesia (n = 3-9 trial blocks; 2-4 mice per condition).

In lobule Crus I, each sensory modality reliably triggered spiking across the PC ensemble that was time-locked to the presentation of individual stimuli (***Figure 1C***). We found slight differences in peak rates of sensory-triggered Ca^2+^ events across modalities (***Figure 1D***). The vast majority of individual PCs within ensembles were responsive to more than one modality (***Figure 1E***). This result suggests that CFs generally report the occurrence of an external stimulus rather than encode modality type in Crus I. However, simultaneously presenting two modalities had a non-additive effect on the rate of CF-evoked events, indicating that ensemble responsiveness to any single stimulus type was near saturation (***Supplemental Figure 1A and 1B***). Interestingly, there was evidence of partitioned processing of sensation and action within distinct regions of the lateral cerebellum. In lobule Crus II, where PCs are responsive to motor-related CF activity (Gaffield et al., 2016; Gaffield and Christie, 2017), sensory cues failed to alter the rate of dendritic Ca^2+^ events compared to baseline (1.22 ± 0.12 and 1.20 ± 0.14 Hz; prior to and during sensory stimuli, respectively; n = 39 trial blocks, 6 mice; p = 0.93; paired Student’s t-test; ***Figure 1C***). Given that sensory-triggered Ca^2+^ events persisted in PCs of Crus I while animals were under anesthesia-induced paralysis, we rule out movement as the direct cause of CF responses in this region (***Figure 1F***).

Examining isolated CF-evoked Ca^2+^ events in PC dendrites (see Methods) revealed that the responses that occurred during sensory stimuli were distinguishable from spontaneous activity based on the responses’ enhanced size (peak integral 27.5 ± 3.2% larger on average; n = 74 trial blocks, 12 mice; P < 0.001; paired Student’s t-test). This result was consistent across modalities (***Figure 2A***). The amount of enhancement varied slightly with stimulus type (***Figure 2B***), which is in line with previous reports of graded coding of sensory-evoked CF activity by PC dendrites (Najafi et al., 2014a, b). However, the subtle differences in sensory enhancement between modality types further suggest that stimuli produce near-saturating responses. This also indicates a limited bandwidth within which graded enhancement can be encoded. Interestingly, sensory enhancement of Ca^2+^ event size was only weakly correlated with firing responsiveness to the stimulus (***Figure 2C***). Even among cells that did not undergo a time-locked change in dendritic spike frequency to cue presentation, sensory stimuli enhanced CF-evoked Ca^2+^ events. Enhancement in these cells was comparable to that in PCs with high rates of sensory-induced spiking (***Figure 2D***). In summary, lobule-specific CFs broadly encode the occurrence of external sensory cues across modalities, generating graded Ca^2+^ signals in PC dendrites that are larger than spontaneous events.

**Figure 2.**
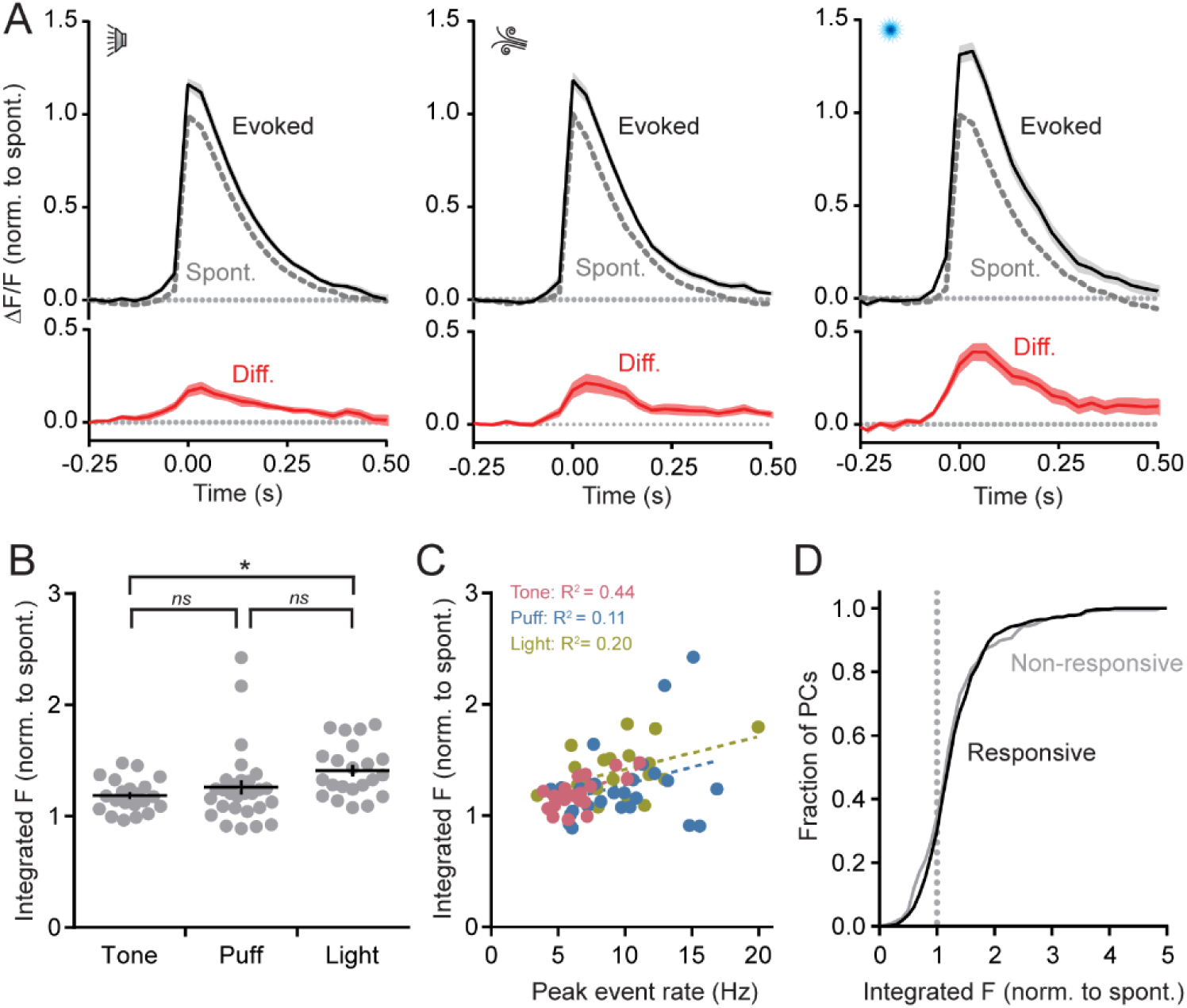
Graded coding of sensory-related Ca^2+^ events in PC dendrites. (**A**)Average isolated dendritic Ca^2+^ events, collected across Crus I PC dendrites from all mice, during sensory stimuli presentation. Peak amplitudes are normalized to that of spontaneous events with the difference plotted in red below. (**B**)The peak amplitude of the integral for average, sensory-associated Ca^2+^ events, relative to that of spontaneous responses, for each block of stimulus trials (indicated by gray dots). Data are mean ± SEM; * indicates p = 0.02, ns: not significant p = 0.59 and 0.12; ANOVA with Tukey’s multiple comparison test (n = 21-28 trial blocks; 6-10 mice per condition). (**C**)Relationship between size enhancement of sensory-related Ca^2+^ events, relative to spontaneous responses, and the induced frequency of dendritic spiking during the stimulus. Each point is the average of a block of trials (6-10 mice per condition). Data were fit with linear functions. (**D**)Cumulative distribution of Ca^2+^ event sizes (integrals normalized to spontaneous responses) from PCs classified as responsive or non-responsive to sensory cue presentation. Responsiveness is based on a peak rate of dendritic spiking during the stimulus that is > 3 standard deviations of the mean spontaneous rate (n = 613 cells from 10 mice).

### Inhibition does not account for differences in CF-evoked dendritic Ca^2+^ signaling

Inhibition from molecular layer interneurons (MLIs) can influence the integrated PC response to CF excitation, reducing the amplitude of dendritic Ca^2+^ signals (Callaway et al., 1995; Kitamura and Hausser, 2011). Thus, graded coding of sensory-triggered CF responses by PCs may, in part, owe to the activity of MLIs. However, convergent lines of evidence argue against this possibility. First, we observed little spontaneous Ca^2+^ activity in RCaMP2-expressing MLIs as animals sat in quiescence (***Supplemental Figure 2A***). This indicates a general lack of inhibitory activity in the absence of behavioral context. To examine for the co-occurrence of MLI activity with spontaneous CF-evoked responses in PCs at high resolution, we averaged dendritic events in GCaMP6f-expressing PCs and the simultaneously measured, corresponding fluorescence signals in surrounding MLIs (***Supplemental Figure 2B***). The absence of a coincident increase in MLI Ca^2+^ activity, time-locked to the PC response, further indicates that inhibition does not suppress the size of spontaneous dendritic Ca^2+^ events in PCs.

Second, MLIs in Crus I were highly responsive to sensory stimuli, with each modality generating robust and widespread activation of their ensemble (***Supplemental Figure 2C and 2D***). Thus, sensory-enhanced PC Ca^2+^ signaling occurred despite the coincident activation of CFs and MLIs during cue presentation, as measured by dual-color Ca^2+^ sensor imaging (***Supplemental Figure 2E***). Surprisingly, the trials that produced the largest changes in PC Ca^2+^ event size were correlated with the greatest levels of MLI activation (***Supplemental Figure 2F***). We would have expected an inverse relationship if MLI-mediated inhibition were directly responsible for producing variable amplitude CF-evoked Ca^2+^ events during cue presentation.Thus, we conclude that a sensory-dependent enhancement of dendritic Ca^2+^ signaling in PCs is encoded through a different neural circuit pathway.

### Sensory enhancement is represented in the presynaptic activity level of CFs

If conveyed to the cortex by CF axons, burst firing in olivary projection neurons (Crill, 1970) could generate graded levels of excitation in postsynaptic PCs dependent on the number of action potentials in the presynaptic burst (Mathy et al., 2009). Sensory enhancement of burst duration could thus contribute to larger dendritic Ca^2+^ responses. Therefore, to directly measure CF activity, we virally transduced olivary projection neurons with GCaMP6f to express this Ca^2+^ indicator in their axons (***Figure 3A***). We readily observed spontaneous Ca^2+^ events in Crus I CFs with an event frequency that closely matched the rate of Ca^2+^ events in PC dendrites, as measured in separate mice (1.04 and 1.01 Hz, respectively; n = 5 and 12 mice; p = 0.75; Student’s t-test). CFs were also activated in response to sensory stimuli with evoked Ca^2+^ event rates that were comparable to the rates observed in PC dendrites (***Figure 3B***). In fact, the fraction of sensory-responsive CFs was identical to that of PCs (***Figure 3C***). These results strongly argue that CFs reliably excite PCs *in vivo*. Moreover, most, if not all, algorithmically identified, dendrite-wide Ca^2+^ events in PCs must be elicited by the activity of CFs.

**Figure 3.**
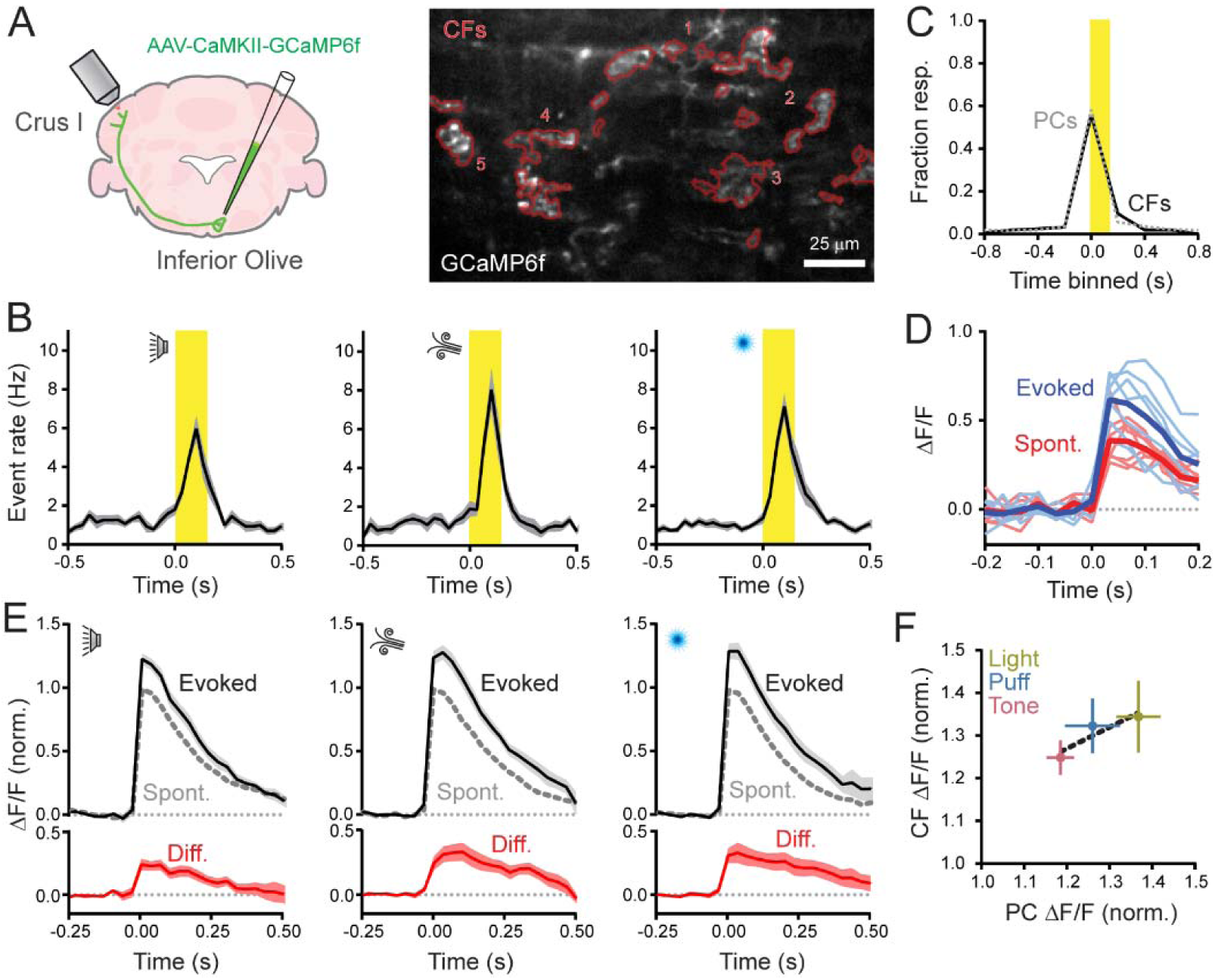
Sensory enhancement of presynaptic Ca^2+^ events in CFs. **(A)**Viral infection of the right inferior olive drove GCaMP6f expression in CFs innervating left Crus I. In the fluorescence image, contours of GCaMP6f-expressing CFs (n = 5) are in red. **(B)**Plots of Ca^2+^ event rates in CFs, evoked by the different sensory modalities. Stimulus timing in yellow (n = 12-15 trial blocks; 5 mice per condition). **(C)**Fraction of sensory-responsive CFs was similar to that of PCs. Measurements were obtained from separate cohorts of animals (n = 613 PCs and 79 CFs; 5 and 10 mice, respectively). **(D)**Isolated Ca^2+^ events, collected from a single CF, were categorized by whether they occurred spontaneously or in response to the sensory stimulus (thick lines are means). **(E)**Average spontaneous and sensory-evoked Ca^2+^ events across CFs for the different sensory modalities; the difference is shown in red (n = 13-16 trial blocks, 5 mice). **(F)**Comparison of the sensory enhancement of Ca^2+^ event amplitudes in CFs compared to observations in PCs, measured in separate experiments. The change in PC response size closely parallels that of CFs across modalities. Data are mean ± SEM.

Next, we examined the size of isolated Ca^2+^ events in CFs and found that responses during cue presentation were larger than responses that occurred outside of the stimulus window (***Figure 3D***). Thus, on average, sensory stimuli produced an enhancement of presynaptic signaling (peak integral 30.9 ± 3.9% larger than spontaneous events; n = 38 trial blocks; 5 mice; p < 0.001; paired Student’s t-test). This result held true across sensory modalities (***Figure 3E***). Notably, there was a strong correlation between sensory-induced enhancement of CF Ca^2+^ event size and the induced increase measured in PC dendrites for each stimulus type (R^2^= 0.84; ***Figure 3F***). In conclusion, the presynaptic activity level of CFs influences the PC dendritic Ca^2+^ response to sensory stimuli.

### Lack of residual dendrite-wide Ca^2+^ signaling in the absence of CF activity

Parallel fiber-mediated excitation of PC dendrites can induce intracellular Ca^2+^ elevation through electrogenic and second-messenger pathways (Ly et al., 2016; Rancz and Hausser, 2006; Tempia et al., 2001). Granule cell activity, driven by sensorimotor stimuli (Chadderton et al., 2004; Chen et al., 2017), could thus directly contribute to the sensory-evoked enhancement of Ca^2+^ signaling in PCs (Najafi et al., 2014b). Therefore, we suppressed the activity of CFs during sensory cue presentation and examined residual, dendrite-wide Ca^2+^ signals in PCs. To accomplish this, we expressed the inhibitory anion-fluxing channelrhodopsin variant *Gt*ACR2 (Govorunova et al., 2015) in excitatory olivary neurons with an adeno-associated virus (AAV) under control of the αCaMKII promoter. Using an implanted optical fiber that targeted the inferior olive, we photoinhibited the activity of *Gt*ACR-expressing cells and imaged Ca^2+^ activity in PCs of Crus I, transduced with GCaMP6f using an AAV under control of the *L7/Pcp2* promoter (***Figure 4A***). During awake quiescence, the frequency of spontaneous dendritic Ca^2+^ events in PCs was markedly reduced during the illumination period (1.15 ± 0.21 and 0.21 ± 0.08 Hz; control and with photoinhibition, respectively; n = 10 epochs, 4 mice; p = 0.005, Students t-test). Ca^2+^ event rates returned to baseline levels after cessation of the optogenetic stimulus (1.30 ± 0.20 Hz). Further analysis revealed that a subset of PCs, comprising 47.9 ± 4.2% of the ensemble population, was significantly affected by optogenetic suppression of the inferior olive.

**Figure 4.**
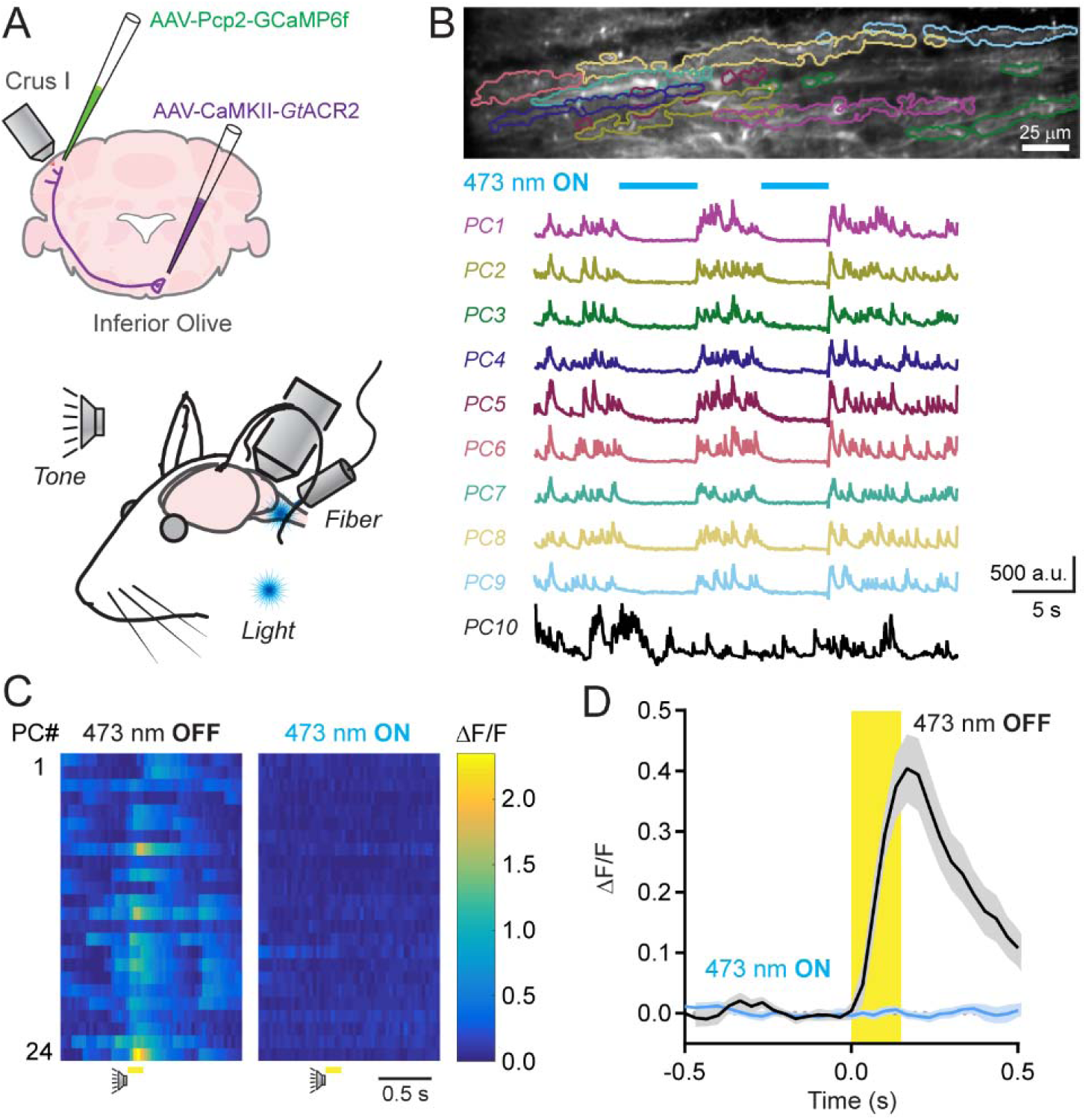
Dendrite-wide Ca^2+^ signals in PCs are completely mediated by CFs. **(A)***Gt*ACR2 was expressed in excitatory neurons of the inferior olive; dendritic Ca^2+^ activity was imaged using 2pLSM in GCaMP6f-expressing PCs of Crus I during sensory stimuli. In some trials, sensory cues were presented to the animal while an optical fiber implant continuously delivered laser light to the inferior olive. **(B)**Spontaneous Ca^2+^ activity in PCs, demarcated in the fluorescence image above, was abolished during illumination of the inferior olive (λ = 473; 250 μW). PC10 illustrates an example PC where activity persisted during illumination. **(C)**Ca^2+^ activity measurements for individual PCs in a field of view in response to an auditory stimulus in control or during the continuous optogenetic suppression of the inferior olive (l = 473; 250 μW). **(D)**Average Ca^2+^ activity transients across all PC dendrites to sensory stimuli (including both auditory and visual cues) in control and with optogenetic suppression of the inferior olive. Data are mean ± SEM (n = 234 cells from 4 mice; 10 trial blocks for each condition).

In these cells, spontaneous Ca^2+^ activity was eliminated (***Figure 4B***). The lack of an effect in the remaining PC population likely reflects incomplete transduction of all olivary projection neurons. Together, these results confirm that spontaneous dendritic Ca^2+^ events in PCs are driven by the output of CFs.

Among PCs that were sensitive to spontaneous activity suppression, optogenetic inhibition of the inferior olive also affected their Ca^2+^ responses to sensory stimulation. When presented with external cues during the continuous photo-suppression of the inferior olive, there was an absence of algorithmically identifiable dendritic Ca^2+^ events within the cue window (1.9 ± 1.3% of control rate; p = 0.002; n = 10 trials, 4 mice; paired Student’s t-test; ***Figure 4C***). Furthermore, in trial-averaged responses aligned to the timing of the sensory stimulus, there was a lack of resolvable dendrite-wide Ca^2+^ signal in PCs during cue presentation (***Figure 4D***).Together, these results indicate that CF activity fully accounts for the sensory enhancement of Ca^2+^ responses in PCs. We conclude that behaviorally relevant information is transferred in a graded manner, at high fidelity, from the inferior olive to PC dendrites.

## Discussion

### PC dendrites broadly encode the occurrence of sensory stimuli across modalities

CF inputs map a fractured somatotopy onto the cerebellar cortex, producing zones of like-responding PCs to sensorimotor stimuli (Apps and Hawkes, 2009). Here, we report lobule-specific responsiveness of PCs to CF-mediated excitation, with sensory stimuli reliably driving activation of PCs in Crus I but not Crus II. CF responses in Crus I were not due to motor action as sensory-related activity persisted under anesthesia-induced paralysis. In previous studies, we also observed lobule-specific differences in the representation of behavioral variables, with orofacial movements in response to reward consumption engaging neurons in Crus II but not Crus I (Gaffield et al., 2016; Gaffield and Christie, 2017). Combined, these results suggest partitioned representations of unpredicted sensations and mechanics of action within the lateral posterior hemispheres, cerebellar regions increasingly implicated in cognitive function in both humans and mice (Deverett et al., 2018; Schmahmann, 2018; Stoodley et al., 2017).Interestingly, we found that individual PCs in Crus I were broadly responsive to multiple sensory modalities. The general lack of CF tuning argues that olivary neurons report the occurrence of a cue rather than a particular sensory event type. This may be conducive for encoding associations to any unexpected external event encountered in the environment and its repercussions to the animal’s behavior.

### PC dendrites integrate the level of presynaptic CF activity

The integrative properties of PC dendrites allow for a large repertoire of CF-mediated responses beyond that of “all-or-none” excitation (Najafi and Medina, 2013). A major finding of our work is that PCs are sensitive to the activity level of CFs, generating graded dendritic Ca^2+^ signals that, on average, change in proportion to size alterations of evoked, presynaptic Ca^2+^ responses. Importantly, enhanced CF signaling occurred in response to external cues. This indicates that behaviorally relevant stimuli, such as an unexpected sensory event, can be differentially encoded by CFs perhaps adding to the saliency of these cues. Because olivary neurons fire variable-duration bursts of action potentials that are transmitted along their axon (Crill, 1970; Mathy et al., 2009), we surmise that presynaptic Ca^2+^ events in CFs correspond to discrete bursts and that the size of these presynaptic Ca^2+^ events reports burst duration. The spike content of olivary bursts changes with the amplitude and phase of subthreshold oscillatory activity in these neurons (Chorev et al., 2007; Leznik and Llinas, 2005). Network coherence and phase-resetting of subthreshold oscillations in olivary neuron ensembles (De Gruijl et al., 2012; Khosrovani et al., 2007) may be influenced by sensory stimuli, promoting amplification of burst duration when animals are confronted with unexpected cues. In turn, sensory amplification of olivary output would be relayed to PCs by CFs and, based on our observations, linearly transformed into dendritic Ca^2+^ signals through locally generated electrogenic potentials (Davie et al., 2008; Otsu et al., 2014).

### Dendritic-wide Ca^2+^signaling in PC dendrites is exclusively mediated by the activity of CFs

In addition to CF-mediated input, PC dendrites integrate excitation and inhibition from parallel fibers and MLIs, respectively, whose activity can modulate the size of CF-evoked Ca^2+^ responses. However, differences in the size of spontaneous and sensory-related, dendrite-wide Ca^2+^ events are probably not attributable to the direct activity of either of these inputs for two reasons. First, MLIs appeared to be relatively inactive during quiescence suggesting that inhibition of CF-evoked dendritic Ca^2+^ signaling in PCs (Callaway et al., 1995; Rowan et al., 2018) does not account for the reduced size of spontaneous events. In fact, in Crus II, chemogenetic disinhibition of the molecular layer has no effect on spontaneous Ca^2+^ event amplitudes (Gaffield et al., 2018). Interestingly, we observed that MLIs in Crus I were co-activated with PCs in response to sensory cue presentation. Thus, although inhibitory output increased, CFs were still able to drive larger dendritic Ca^2+^ events in this behavioral context. Second, we failed to observe residual Ca^2+^ signal in PC dendrites in response to sensory stimulation when the output of the inferior olive was optogenetically suppressed. If sensorimotor-related parallel fiber activity directly triggered a dendrite-wide Ca^2+^ signal (Najafi et al., 2014b), then this should have been revealed in this condition.

Our results do not rule out possible roles for these modulatory inputs in graded coding of behaviorally relevant PC Ca^2+^ signals. For example, the coincident activity of parallel fibers can produce an indirect, non-linear enhancement of CF-evoked responses, apart from direct Ca^2+^ entry, by affecting PC dendritic excitability (Otsu et al., 2014; Wang et al., 2000). However, this non-linear enhancement is tightly regulated by feedforward inhibition (Gaffield et al., 2018), and any behavioral contexts that enable granule cells to influence CF-evoked responses have yet to be resolved. It is also possible that parallel fibers and/or MLIs have a subcellular influence on PC dendritic Ca^2+^ signaling (Callaway et al., 1995; Roome and Kuhn, 2018), not apparent in the average, arbor-wide response. We hypothesize that sensory enhancement of CF burst firing may aid in the efficacy of learning (Najafi and Medina, 2013). By generating a corresponding increase in the size of PC dendritic Ca^2+^ signals, longer duration CF bursts can help determine the direction and strength of short- and long-term synaptic plasticity at parallel fiber inputs (Mathy et al., 2009). In the absence of a dendrite-wide modulatory influence through the mossy fiber pathway, information is transferred from CFs to PCs at high fidelity, ensuring a reliable coding scheme whereby instructive signals for plasticity and learning are entirely conveyed from the inferior olive to the cortex.

## Methods

### Mice

Our experiments were conducted under approval of the Institutional Animal Care and Use Committee at the Max Planck Florida Institute for Neuroscience. Adult mice (>10 weeks of age) of both genders (13 female, 11 male) were used for all experiments. We used heterozygous *Kit::Cre* mice (n = 16; Amat et al., 2017), homozygous *Pcp2::Cre* (n = 4; Barski et al., 2000; Jax #004146), or C57BL/6J mice (n = 4; Jax #000664). All mice were housed in reverse light/dark cycle and received enrichment with running disks.

### Surgical Procedures

Surgeries were performed under isoflurane anesthesia (1.5-2.0%). Once a deep plane of anesthesia was induced, determined by the lack of response to a toe pinch, a lubricating ointment was applied to the eyes and the animal was injected subcutaneously with carprofen (5 mg/kg), dexamethasone (3 mg/kg), and buprenorphine (0.35 mg/kg) to reduce post-surgical swelling and pain. The hair covering the top of the skull was removed and the exposed skin disinfected with iterative scrubs of iodine and ethanol solutions. Throughout surgery, body temperature was maintained at 37°C using a heating pad with biofeedback control. Topical application of lidocaine/bupivacaine to the surgical site provided local anesthesia. The skin overlaying the skull was excised, and then the skull was gently scraped clean with a scalpel before a custom-made stainless steel head post was attached using dental cement (Metabond; Parkell, Edgewood, NY). A small craniotomy about 2 mm square was cut over left CrusI/II centered approximately 3.5 mm laterally and 2.2 mm caudally from lambda. This craniotomy exposed portions of zebrin bands 7+, 6-, and 6+ of both lobules. Care was taken to avoid damaging the dura.

AAVs, singly or in combination, were injected through a lone, beveled-tip glass micropipette into the exposed brain. AAVs injected into this area of cortex included the following (acquired from the University of North Carolina Vector Core Facility, the University of Pennsylvania Vector Core Facility, or ViGene, Rockville, MD): PCs were targeted in *Pcp2::Cre* mice using AAV1-CAG-Flex(*loxP*)-GCaMP6f. In *Kit::Cre* mice or C57BL/6J mice, PCs were targeted using either AAV1-Pcp2.4-CMV-GCaMP6f (Nitta et al., 2017) or AAV1-Pcp2.6-FLPo (Nitta et al., 2017) in combination with AAV1-CAG-Flex(FRT)-GCaMP6f at a 1:1 ratio. MLIs were targeted in *Kit::Cre* mice using AAV1-CAG-Flex(*loxP*)-RCaMP2 either alone or in combination with AAV1-Pcp2.6-FLPo and AAV1-CAG-Flex(FRT)-GCaMP6f at a 1:1:3 ratio. A total volume of 250-350 nL of high titer viral particles (e^12–e^13) were pressure injected into the tissue at a rate of ~25 nL/min. Intraperitoneal injections of *D*-mannitol (750 mg/kg) promoted viral spread. For targeting the inferior olive, viral particles (AAV1-αCaMKII-GCaMP6f, AAV1-αCaMKII-*Gt*ACR2-eFYP or, in one animal, AAV1-αCaMKII-*Gt*ACR2-ST-FRed-Kv2.1 to restrict expression to the soma) were injected into the brainstem (~500 nL; 25 nL/min) at the following coordinates: x = 0.3 mm, y = −4.9 mm, z = −4.6 mm, at a depth of 3.6 mm and an approach angle of 62°. In optogenetic experiments, an optical fiber (CFMLC21U; 105 μm, 0.22 NA, 3.5 mm in length; Thorlabs; Newton, NJ) was then inserted using the same coordinates, and secured in place with dental cement.

After completion of viral injections, a small glass coverslip (CS-3R; Warner Instruments, Hamden, CT) was attached to the skull over the craniotomy using cyanoacrylate glue and then encased with dental cement that also covered any exposed parts of the skull. Care was taken to minimize the amount of pressure exerted by the glass onto the brain’s surface. The animals were recovered until ambulatory and monitored for signs of stress and discomfort for 7 days before proceeding to further manipulations.

### Two-photon Microscopy and Optogenetics

2pLSM was used for measuring *in vivo* Ca^2+^ activity in neurons of the lateral cerebellum with a custom-built, movable-objective microscope as described previously (Gaffield et al., 2016; Gaffield and Christie, 2017). Video-rate frame scanning (~30 frames/s) at high resolution (512 × 512 pixels) was achieved using an 8 kHz resonant scan mirror in combination with a galvanometer mirror (Cambridge Technologies; Bedford, MA). We used a 16x, 0.8 NA water immersion objective (Olympus; Center Valley, PA) and diluted ultrasound gel (1:10 with distilled water) as an immersion media (Aquasonic; Parker Labs, Fairfield, NJ). The two-channel light-collection pathway, separating green and red light, used high-sensitivity photomultiplier tubes for detection of emitted photons (H10770PA-40, Hamamatsu; Bridgewater, NJ). The microscope was controlled using ScanImage 2015 software (Vidrio Technologies; Ashburn, VA). GCaMP6f (Chen et al., 2013) was excited at 900 nm (Chameleon Vision S, Coherent, Santa Clara, CA). For PCs, we used <50 mW of power at the objective and ~100 mW for imaging CFs. RCaMP2 (Inoue et al., 2015) was excited at 1070 nm (Fidelity 2, Coherent) with <60 mW of power.

Optogenetic stimulation of *Gt*ACR2 was driven by 473 nm light from a continuous-wave laser (MBL-F-473-200mW; CNI Optoelectronics; Changchun, China). The laser path included an acousto-optic modulator for precise control of light timing (MTS110-A3-VIS controlled by a MODA110 Fixed Frequency Driver; AA Opto-Electronic; Orsay, France). Laser light was launched into a fiber port (PAF-X-11-A; Thorlabs) and directed into a patch cable (105 mm diameter core, 0.22 NA; M61L01; Thorlabs). This delivered laser light to the optically matched fiber implant. A ceramic connector (ADAL1; Thorlabs), covered with two layers of shrink tubing and masking tape, limited escape of stray light and secured the patch cable to the implant. Light out of the patch cable was <1 mW.

### Behavior and Sensory Stimuli

Mice were head restrained by their surgically attached head post during neural activity monitoring; prior acclimatization allowed familiarity to restraint and improved the stability of recordings. Likewise, animals were water restricted (1 mL/day) to associate restraint to reward allocation, a procedure that improved recording stability. All water was provided before beginning any trials.

For sensory stimuli, a pure auditory tone (12 kHz) was generated by a microcontroller (Mega; Arduino; Ivrea, Italy) attached to a speaker positioned on the right side of the animal (~85 dB at the animal’s ear). Somatosensory vibrissal stimulation was achieved using a focused airpuff (18 psi; from a compressed nitrogen source), delivered from a needle positioned 3 cm from the face, onto the left whisker pad. A visual stimulus was generated by a 473 nm light source (either a laser or LED), transmitted through the capped end of an optical patch cable (25 μW) visible from the left eye. All stimuli were 150 ms in duration. The same stimulus type was repeatedly delivered (15x) in a block of trials (0.5 Hz). For each block of trials, the order of each modality was randomized to limit habituation. Stimulus presentation as well as task timing and monitoring were controlled with bControl software (Brody Lab; Princeton). Optogenetic suppression trials were interleaved with control trials to confirm responsiveness to sensory stimuli. For experiments conducted under anesthesia, ketamine/xylazine (100 and 10 mg/kg, respectively) was delivered by intraperitoneal injection.

### Data Analysis

All data were analyzed using custom-written routines in MATLAB (MathWorks; Natick, MA). Time-series images of Ca^2+^ activity in neurons expressing genetically encoded Ca^2+^ indicators were aligned using a least-squares algorithm. PCs and CFs were segmented using an independent component analysis algorithm to group like-responding pixels (Hyvarinen, 1999) and Ca^2+^ events identified using an inference-based method (Gaffield et al., 2016; Vogelstein et al., 2010). Ca^2+^ event rates were calculated from each image (34 ms in duration). Cell responsiveness to sensory stimuli was determined if the number of detected events increased > 3 standard deviations from the baseline event rate.

For analysis of Ca^2+^ event size, we focused on discrete, isolated responses (Gaffield et al., 2018). Thus, we avoided potential uncertainties associated with GCaMP6f non-linearity (Chen et al., 2013) for overlapping events. For inclusion, events must have been separated by at least 500 ms from proceeding or following events for PCs and 300 ms for CFs, a time window allowing for a nearly complete decay of the response. Events were categorized as sensory-related if they occurred within 250 ms of stimulus onset. Events outside of this window were categorized as spontaneous. To quantify event size, we measured the peak of the integral of the Ca^2+^ response, a metric identical to previous reports (Najafi et al., 2014a, b). ΔF/F was calculated using a baseline fluorescence period immediately prior to an identified event (200 ms). Lastly, for trial-averaged PC Ca^2+^ activity measurements that were irrespective of isolated events, we simply calculated ΔF/F for all PC dendrite regions of interest (ROIs) and aligned responses to the onset of the sensory stimulus.

MLI ROIs were hand-drawn from time-averaged images; automated segmentation was difficult because MLIs respond in a nearly identical manner (Gaffield and Christie, 2017). Therefore, for population analysis, we used an ROI encompassing the entire field of view to measure and quantify sensory-evoked Ca^2+^ activity. For dual-color Ca^2+^ imaging, epochs of spontaneous MLI activity were temporally registered to that of CF-evoked Ca^2+^ events in PCs; a mask comprising all pixels corresponding to MLIs within 50 μm of each PC was used to calculate the time-locked average of MLI activity during these periods.

Additional calculations and plotting was performed with Excel (Microsoft, Redmond, WA) and Prism (GraphPad, La Jolla, CA). Statistical analysis was performed in Excel or Prism. Significance was defined as having a p value < 0.05. Student’s t-tests and ANOVAs were used where appropriate as indicated in the text. In figures, neural activity traces are shown with the mean response in bold and ± SEM in the shaded region. In summary plots, data are represented as mean with error bars indicating SEM. We did not perform a power analysis prior to experiments to estimate replicate numbers.

## Acknowledgements

We thank the GENIE program (Janelia Research Campus, including Drs. Jayaraman, Kerr, Kim, Looger, and Svoboda) for freely providing GCaMP6f to the neuroscience community, Dr. Bito (University of Tokyo) for use of RCaMP2 and Dr. Yizhar (Weizmann Institute of Science) for use of somatically targeted *Gt*ACR2. This work was supported by the Max Planck Society, the Max Planck Florida Institute for Neuroscience, and National Institutes of Health Grants NS083894 and NS105958 (J.M.C).

## Author Contributions

Conceptualization, M.A.G. and J.M.C.; Methodology, M.A.G.; Investigation, M.A.G.; Writing, M.A.G. and J.M.C.; Funding Acquisition, J.M.C.

### Declaration of Interests

The authors declare no competing interests.

---

**Supplemental Figure 1.**
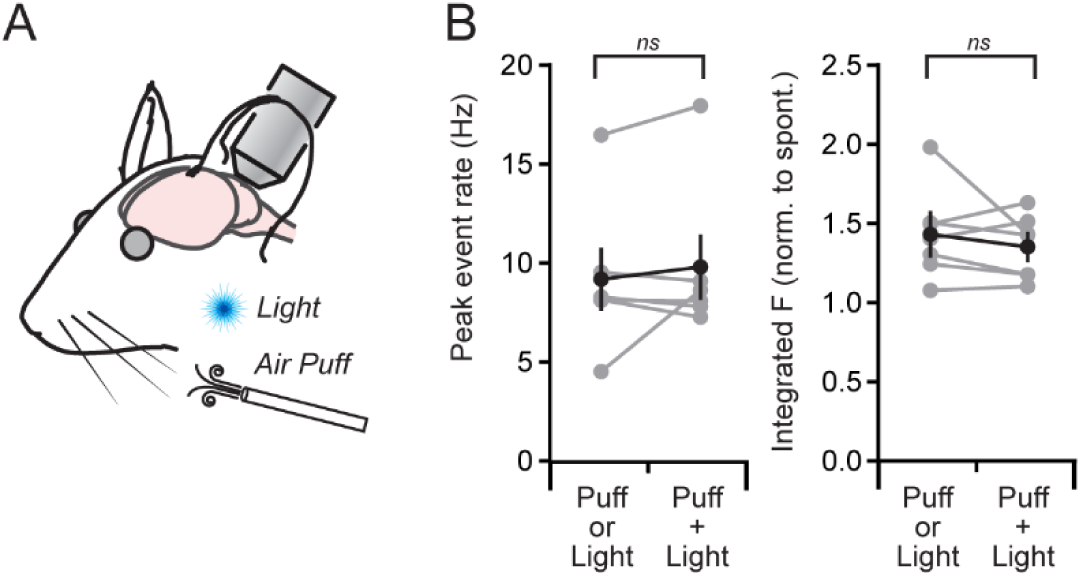
Non-additive effect of simultaneous presentation of different sensory stimuli on PC Ca^2+^ events. (**A**)In the same mouse, a visual or a somatosensory stimulus was presented alone or in simultaneous combination. (**B**)Summary plots showing the change in frequency and size of PC dendritic Ca^2+^ events both to individual sensory modalities or to their simultaneous presentation. Data are mean ± SEM with individual mice in gray; ns, not significant, p = 0.45 and 0.40; paired Student’s t-test (n = 6-7 mice).

**Supplemental Figure 2.**
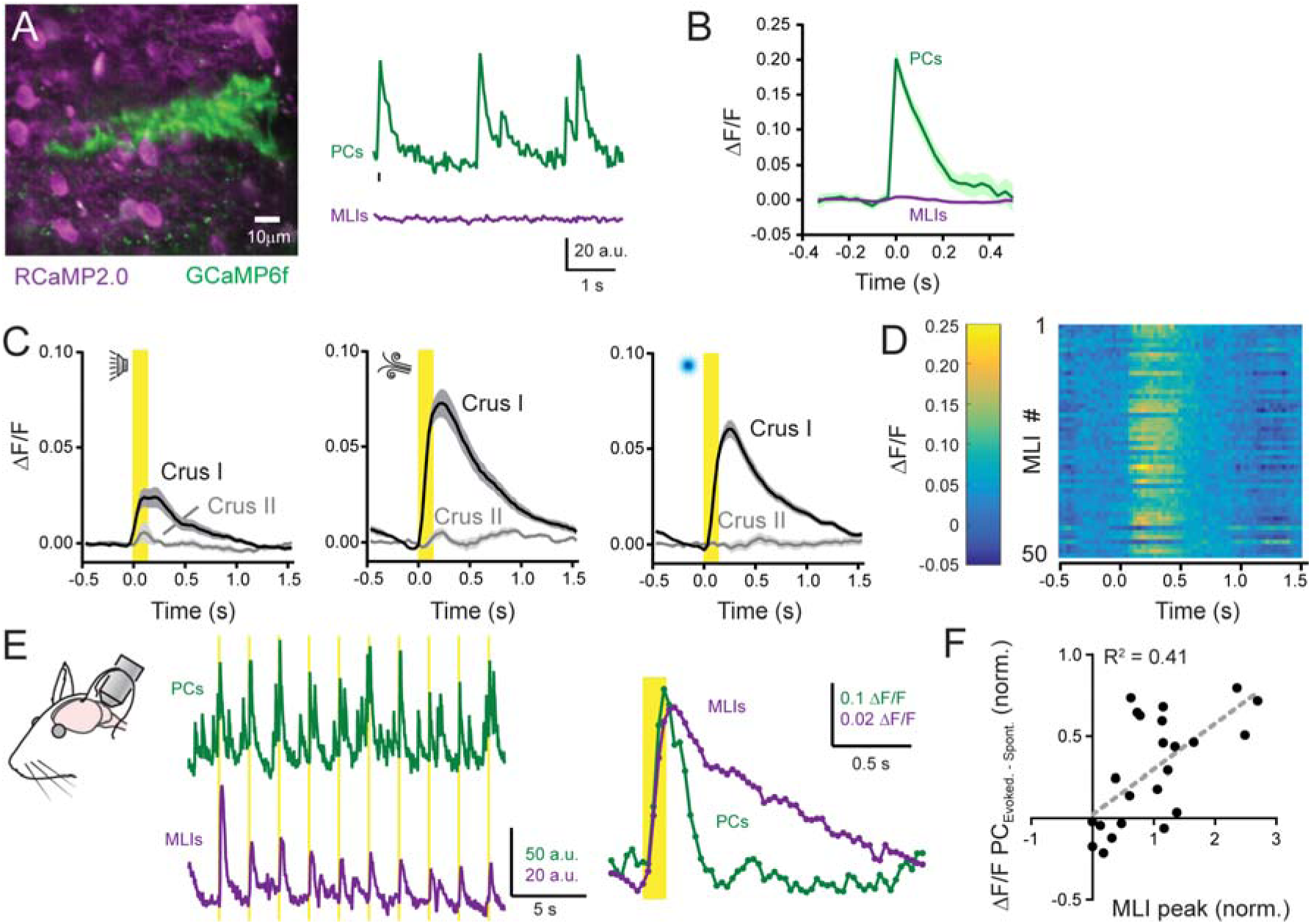
MLIs do not account for differences in sensory-evoked Ca^2+^ events in PC dendrites. (**A**)Left: Image of an isolated, GCaMP6f-expressing PC dendrite surrounded by RCaMP2-expressing MLIs. Right: Simultaneous, dual-color Ca^2+^ activity measurements from PCs and MLIs during quiescence. An isolated dendritic Ca^2+^ event is demarcated by the black tic mark. (**B**)Average of MLI Ca^2+^ activity aligned to the occurrence of identified spontaneous CF-evoked dendritic Ca^2+^ events (n = 212; 3 mice) in nearby PCs (within 50 μm). **(C)**Trial-averaged ensemble responses of MLIs to different sensory modalities, measured in two different lobules across mice (n = 6-20 trial blocks; 5-7 mice per condition). **(D)**Ca^2+^ activity measurements from all identified MLIs in a single field of view from one mouse. The responses are an average of the repeated stimulation of whiskers with an air puff (n = 15). **(E)**Left: Simultaneous dual-color activity measurements from PCs and MLIs during repeated stimulation of the whiskers with an air puff. Yellow bars indicate timing of the sensory stimuli. Right: Superimposition of sensory-evoked Ca^2+^ events in PC dendrites and the response elicited in the surrounding MLI ensemble. **(F)**Relationship between the cue-dependent enhancement of Ca^2+^ events in PC dendrites and the peak amplitude of the sensory-evoked response in MLIs. Each point represents the average response from simultaneous activity measurements of MLI ensemble activity and the change in PC dendritic Ca^2+^ size from a block of trials (measurements from 3 mice).

